# Semisynthetic ferritin nanocages for flexible, site-specific targeting, cluster-formation and activation of membrane receptors

**DOI:** 10.1101/2024.11.01.621585

**Authors:** Andreas Neusch, Christina Siepe, Liesa Zitzke, Alexandra C. Fux, Cornelia Monzel

## Abstract

Homopolymerization and cluster formation of cellular membrane receptors (MR) is closely related to their signaling activity. However, underlying mechanisms and effects of clustering are often hardly understood. This lack of knowledge is due to the lack of suitable tools which enable to specifically target and activate distinct MRs, without causing side-effects. In this study, we designed a fluorescent semisynthetic nanoparticle (NP) based on the iron-storage protein ferritin and *S. aureus* Protein A, that is readily equipped with a variety of antibodies with *K*_D_ values below 5 nM. Specificity of the NP antigen recognition was evaluated in cell experiments with cells expressing Transferrin Receptor 1 or the death receptor CD95, both of which displayed rapid cluster formation upon contact with the NP. Lastly, it was possible to induce apoptosis solely by induced clustering of CD95 via our engineered NP.

## MAIN TEXT

Living cells are constantly facing multitudes of incoming information through all sorts of extracellular molecules. In order to process these signals effectively, intricate signaling pathways have developed around membrane receptors (MR) that are specifically recognizing distinct extracellular molecules, so called ligands.^1^ To this date, more than 1350 MRs were identified in the human proteome.^2^ Upon ligand recognition, MRs initiate signaling pathways, that can involve complex, multilayered chemical cascades that ultimately trigger a cellular reaction. Malfunction of signaling pathways was shown to be closely related to various diseases.^3–6^

Signaling pathways are often initiated by homopolymerization of MRs. In this context, the role of supramolecular structures and spatial arrangements of receptors involved in signaling has been highlighted in several studies.^7–12^ However, the exact mechanisms behind the formation of these clusters and the subsequent signaling are largely unknown. This might partly be because of the lack of molecular tools that can trigger distinct MRs specifically, without affecting the cell through off-target effects.^13,14^ Among the recent approaches to induce MR clustering, photoactivatable reagents,^15,16^ magnetogenetics,^17–19^ and optogenetics,^20,21^ constitute powerful tools to locally control the protein activity of cells and to learn about the signal initiation mechanism. However, these approaches often require genetic modifications of the proteins of interest and the application of sophisticated stimuli.

Among all known MRs, the tumor necrosis receptor family (TNF-R) is considered one of the best known. It is responsible for cell proliferation, survival, immune homeostasis, and the programmed cell death, apoptosis.^11,22^ Cluster of differentiation 95 (CD95) is one death receptor of the TNF-R family, that is able to trigger apoptosis upon binding to its ligand CD95L.^23^ Prior to initiation of apoptosis, some CD95 may form small oligomers^12^ and lipid rafts containing CD95 were shown to promote receptor clustering. After ligand binding CD95 oligomerizes at least to dimers and trimers at characteristic distances and initiates the subsequent apoptosis signal.^24–27^ However, it was also shown that pre-ligand assembly domains (PLADs) are promoting dimerization and signal initiation in TNF-Rs in the absence of ligand.^28^ While excessive apoptosis has been linked to diseases such as Alzheimer’s and Parkinson’s, insufficient apoptosis can lead to excessive cell growth and, thus, cancer.^29,30^ Therefore, tools that are able to specifically trigger and control cell death, are of high medical interest.

Another membrane receptor that is connected to cancer is Transferrin Receptor-1 (TfR1). TfR1 is responsible for cellular iron uptake and iron homeostasis in general, by binding and internalizing the iron transporter Transferrin.^31^ Furthermore, TfR1 has been found to be overexpressed in cancer cells, making it a suitable marker for tumor cells.^32^

Here, we designed a nanoparticle (NP) platform, that is able to site-specifically target membrane-bound receptors. We focused on receptors that were shown to be overexpressed in cancer cells. After receptor targeting, our NP is designed to induce receptor clustering and, if applicable, MR activation.

One highly promising approach for efficient targeting is the ferritin family, an ubiquitous class of iron-storage proteins. Present in nearly all forms of life, ferritins feature a spherical, hollow protein cage that is highly conserved and serves as storage for iron ions.^33,34^ For scientific application, the most commonly used member is human ferritin, with a diameter of 12 nm, while its inner cavity is 8 nm in size.^35^ Here, the protein cage consists of 24 subunits of an arbitrary combination of either L chain ferritin (LCF) or H chain ferritin (HCF),^36,37^ named after the locations where the subunits were first isolated from: liver and heart.^38^ Coincidentally, LCF (19 kDa) is also slightly lighter than HCF (21 kDa), thus the two subunits are often referred to as light and heavy chain. The main difference between the two subunits is HCF’s ferroxidase domain, that oxidizes toxic ferrous iron to insoluble, harmless ferric iron.^39^

There are several advantages of applying ferritin as a platform for MR targeting. As a protein, it can fairly easily be modified through genetic engineering. To this end, its 24-mer cage can be equipped with different tags, in order to create a multi-functional nanoparticle (NP). Due to ferritins human origin, it has low immunogenicity and is, in contrast to many synthetic NPs, biodegradable. Plus, ferritin was shown to be a natural ligand to TfR1.^40^ More specifically, the coupling is mediated by HCF.^41^ Upon binding, TfR1 initiates endocytosis of both itself and ferritin.

In this study, we focused on the development of an NP platform for specific targeting. One feasible approach is to use antibodies as targeting mediators (see Figure 1). Immunglobulin G (IgG) is the most common antibody (Ab) and consists of two light and two heavy chains that combine to form the characteristic Y-shape with a molecular mass of ∼ 150 kDa (see Figure 1B).^42,43^ The specific binding site is formed between a heavy and a light chain at the outer edge of the protein, the so-called fragment antigen-binding (Fab) region. The constant region consists of two heavy chains that form the fragment crystallizable (Fc) region, that serves as a backbone to stabilize the structure.^44^ In order to allow the antibody to retain its antigen-binding capabilities after coupling to an NP, it is crucial to connect the two via the Fc region. This leaves the target binding sites free of steric hindrance and, hence, functional. One way to establish such an oriented connection is through Protein A (SpA) from the bacterium *Staphylococcus aureus*.^45^ This 42 kDa protein consists, among others, of five domains of about 58 amino acids each that form one antibody binding domain.^46^ One of these domains, the B-domain, was isolated and modified to give rise to the Z-domain (from here on named protA) that is capable of binding antibodies on its own with a *K*_D_ value of 20 nM.^47,48^ Genetically fusing this domain to ferritin allows direct coupling of IgG to the protein’s surface via incubation (see Figure 1C). This flexible approach enables a quick and straightforward creation of highly specific NPs, with the only constraint being the need for a compatible antibody.^49^

**Figure 1.**
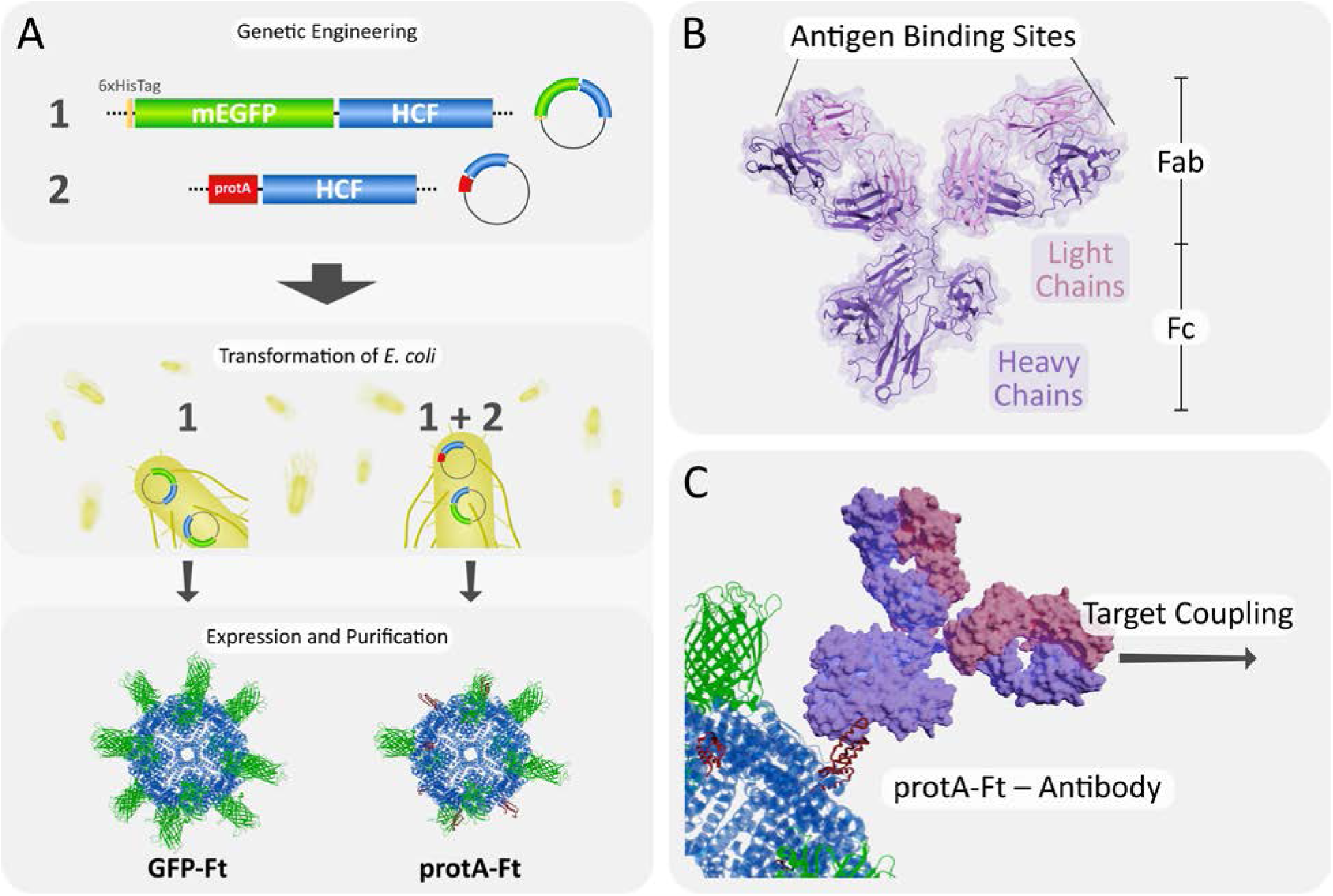
(A) Schematic display of the generation of ferritin nanoagents. Two constructs were genetically engineered: His6-mEGFP-HCF and protA-HCF (see Figure S3 and S4). Plasmid DNA was expressed in E. coli and subsequently purified to obtain both forms, GFP-Ft and the hybrid protA-Ft. (B) Structure of an IgG antibody. (C) Basic principle of protA-Ft activation via antibody coupling. Activated protA-Ft is able to specifically couple to the antibody’s target.

In addition, guiding NPs to their target is as important as detecting and observing them. To this end, we chose to attach a fluorescent probe in the form of monomeric enhanced green fluorescent protein (mEGFP) to our NPs.^50,51^ Using genetic engineering, two ferritin-based constructs were created: (1) mEGFP coupled to HCF (see Figure S3),^52^ and (2) protA coupled to HCF (see Figure S4). Through the nature of ferritin’s self-assembling properties, we produced both a control cage consisting only of mEGFP-HCF (hereafter GFP-Ft) and a hybrid cage combining both constructs, resulting in a mEGFP-protA-ferritin cage (hereafter protA-Ft, see also Figure 1A). This hybrid form is easily trackable via its fluorescent subunits and can also be easily equipped with a matching antibody to direct it to any specific target on – or even in – a cell (see Figure 1C)

The two ferritin constructs were expressed in the *E. coli* strain BL21-CodonPlus (DE3)-RIPL and purified using IMAC. In order to analyze the particles’ physical properties, they were analyzed using transmission electron microscopy (TEM), dynamic light scattering (DLS) and sodium dodecylsulphate polyacrylamide-gel electrophoresis (SDS-PAGE) (see Figure 2 and Table 1). Most notably, all three methods demonstrated the high purity and monodispersity of the ferritin samples. Molecular weights of the monomers matched our expectations. TEM and DLS data revealed cage structures consistent with previously reported (recombinant) ferritins.^52–54^ The observed size difference between TEM and DLS is explained by the experimental setup: while TEM captures the dry protein shell, DLS measures particle properties in solution, hence also including the particle’s hydration shell. Consequently, this leads to a larger detected particle size, namely the hydrodynamic diameter *D*_H_. Furthermore, all data suggests that no significant changes in both cage size as well as cage morphology were introduced by the shift from homo-to heteropolymeric ferritin. The successful creation of heteropolymeric ferritin is evident from the two bands that appeared in SDS-PAGE, representing GFP-HCF and protA-HCF (see Figure 2C).

**Figure 2.**
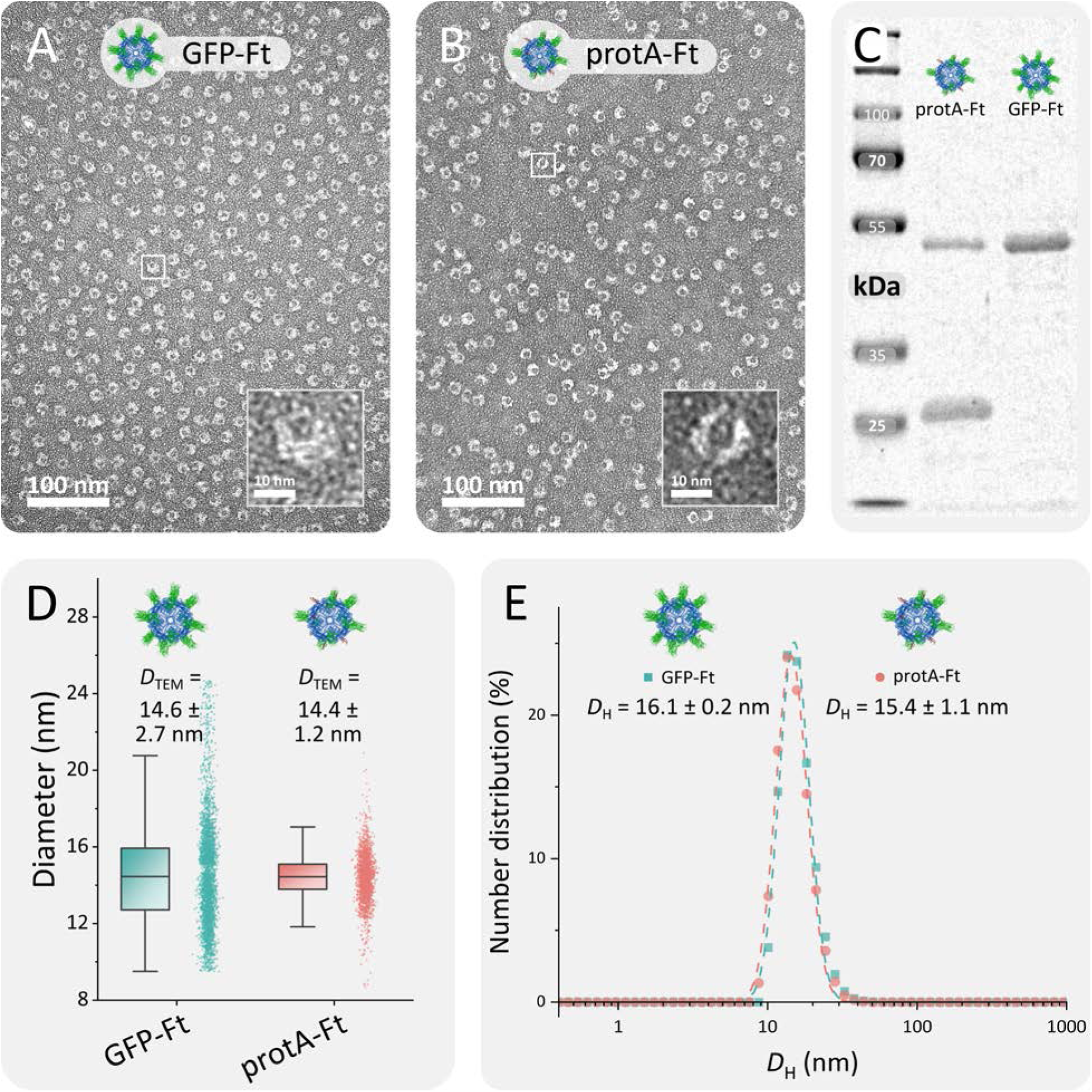
Characterization of ferritin NPs. (A) and (B) show TEM images of GFP-Ft and protA-Ft with an inlet showing an enlarged single cage. (C) SDS-PAGE of both constructs. The first lane shows a protein standard ladder with components of known size (shown in kDa). Background of the image was subtracted using a sliding paraboloid with a radius of 50 pixels. (D) Size distribution of NPs from TEM images. (E) DLS data showing the number distribution of the size of both NP constructs.

**Table 1.** Composition and properties of ferritin constructs used in this study. Since the exact composition of protA-Ft is unkown, the molecular weight can only be estimated. (MW: molecular weight, DTEM: diameter obtained from TEM imaging, DH: hydrodynamic diameter and PdI: Polydispersity index, both from DLS)

| Name | Monomer | MW (kDa) | | $D_{TEM}$ (nm) | $D_H$ (nm) | Pdl |
| --- | --- | --- | --- | --- | --- | --- |
|  |  | Monomer | Cage |  |  |  |
| GFP-Ft | His6-mEGFP-HCF | 49.8 | 1194.7 | $14.6 \pm 2.7$ | $16.1 \pm 0.2$ | $0.13 \pm 0.01$ |
| protA-Ft | His6-mEGFP-HCF | 49.8 | ~900 | $14.4 \pm 1.2$ | $15.4 \pm 1.1$ | $0.11 \pm 0.04$ |
|  | protA-HCF | 28.5 |  |  |  |  |

The two ferritin constructs were further characterized by spectroscopically measuring the degree of labeling (DoL), a measure for the number of fluorescent species coupled to a NP. The DoL of ferritin and mEGFP can be calculated using Beer-Lambert’s law^55^

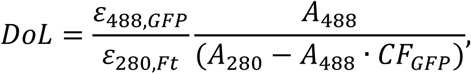

with the absorbance *A*, the molar extinction coefficient *ε*, both at the specified wavelength and the correction factor CF_GFP_ that is given by

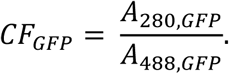

The DoL for GFP-Ft was measured to be 11.2 ± 0.5 GFP/Ft (see Figure S1). While these values strongly deviate from theoretically possible values of up to 24 GFP/Ft, they give an insight into the surface functionalization of our ferritin constructs. It seems, that despite consisting of 24 GFP-labeled HCF subunits, only about 50% of the GFP subunits are indeed actively fluorescent. Explanations for the decreased activity states include (1) steric hindrance on ferritin’s limited surface area, that disturbs GFP structure and hence hampers its activity, (2) post-translational cleavage between the two fused subunits HCF and GFP, (3) HOMO-FRET events that influence absolute absorbance values or (4) intrinsic folding and activity deficits when expressing GFP homologues in *E. coli*.^56^ Nonetheless, a distinct difference was detectable between GFP-Ft and protA-Ft with a DoL of 3.0 ± 0.0 GFP/Ft (see Figure S1). This difference can be seen as an indication that the composition of the heteropolymeric protA-Ft is roughly 1 to 3 (GFP-HCF to protA-HCF). This estimation is supported by SDS-PAGE that shows a ratio of GFP-HCF to protA-HCF of roughly 1 to 2. It seems therefore, that the hybrid protA-Ft cage consists to a notable excess of protA-HCF, giving rise to more than twelve antibody-binding sites.

After the initial characterization, we assessed whether the natural binding capabilities of ferritin towards TfR1 are constrained. In order to precisely monitor coupling and changes of cells coming in contact with protA-Ft, we used a microshower system to incubate single cells with protA-Ft (see Figure 3A). A solution of protA-Ft was loaded into a microcapillary and then placed several µm above the cell of choice while maintaining a constant outflow. This created a stable cloud of ferritin NPs around the cell, similar to localized, cell-specific incubation in NP-loaded media.

**Figure 3.**
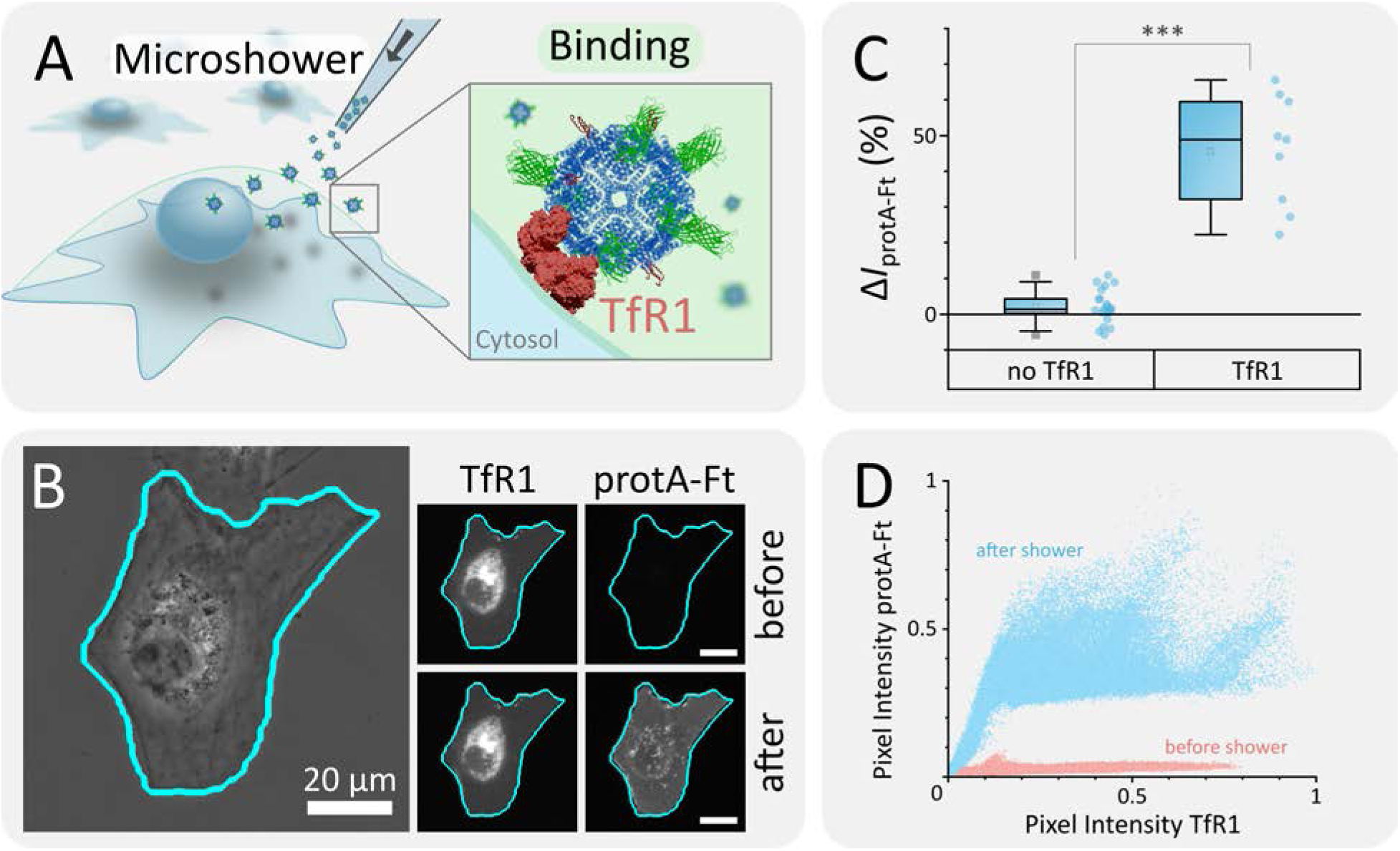
Targeting of TfR1 using protA-Ft. (A) Schematic experimental setup. Cos7 cells were showered with a microcapillary filled with protA-Ft for 15 min. Binding to cells occurs on cells that are overexpressing TfR1. (B) Microscopy data of exemplary cell before and after protA-Ft microshower. Shown are phase contrast image, red channel (TfR1 – pHuji, contrast enhancement of 1 %), and green channel (ferritin, mEGFP). (C) Intensity increase ΔI in protA-Ft’s fluorescence channel after microshower for 15 min in wild type cells (‘no TfR1’) and cells overexpressing TfR1 (‘TfR1’). A one-way ANOVA was performed to test for differences between groups (*** = p < 0.001). (D) Scatter plot of pixel values from TfR1 channel vs. protA-Ft channel, both normalized.

In contrast to wild type cells, protA-Ft clearly bound to the membrane of TfR1-overexpressing cells after a microshower of 15 min (see Figure 3B). Fluorescence intensity in ferritin’s green channel significantly increased by a mean of 46 ± 16 % when showering cells with TfR1, while intensities in wild type cells hardly changed by 2 ± 5 % (Figure 3C). Note that due to the nature of epifluorescent microscopy, intensity values include out-of-focus. Hence, pixel values represent – to some extent – Z-scans across the cell and are therefore brighter towards its thicker center, where the nucleus and endoplasmic reticulum (ER) are located.

Direct comparison of pixel values of the red (pHuji on TfR1) and green (mEGFP on protA-Ft) channels showed that after the microshower, a linear dependency of the two emerged, while *a priori*, those two variables were basically independent (see Figure 3D). We have deliberately not applied Pearson’s Correlation to our microscopic data to avoid misinterpretation due to background signal and cellular autofluorescence in the green channel, but this pixel-wise comparison nonetheless demonstrates high colocalization of TfR1 and protA-Ft. Therefore, we can conclude that despite our implemented modifications, the natural binding capabilities between TfR1 and ferritin are still present. On the other hand, we saw that further modification to the ferritin cage, namely the passivation using polyethylene glycol (PEG), disturb binding to TfR1 and suppresses the correlated intensity increase (data not shown). It is therefore advisable to find an ideal state of modification to not hamper ferritins binding behavior.

After the initial characterizations and testing, we proceeded towards the application of protA-Ft as a targeting hybrid NP. To activate protA-Fts targeting capabilities, antibodies are conjugated to the protA domains on the NPs surface. For characterization of this coupling, a fluorescence-linked immunosorbent assay (FLISA) was performed.^57^ Here, target antibody was immobilized on the wells of a 96-well plate, followed by incubation with protA-Ft. After washing off any excess protein, the fluorescence of protA-Ft remaining inside each well could be measured to evaluate the binding capabilities between protA-Ft and the target antibody. In order to demonstrate the versatility of our hybrid ferritin construct, we characterized the binding of various control antibodies: human IgG1, murine IgG2a and a fluorescently labeled IgG1-APC antibody. As shown in Figure S2, protA-Ft bound to all examined antibodies with *K*_D_ values in the range from 1.6 to 3.7 nM.

This even undercuts previously reported *K*_D_ values for the minimalized protA from *S. aureus* by almost one magnitude.^48^ This decrease can be explained by the avidity effect: protA binds to the antibody’s heavy chain in its Fc region. Since antibodies in general consist of two heavy chains (see also Figure 1), there are two potential binding sites for protA.^58^ Our hybrid protA-Ft cage is equipped with multiple protA subunits. Thus, the antibody is not only bound to one binding site but chelated by two subunits. This in turn entropically strengthens the bond between ferritin and antibody. We observed bonding between protA-Ft and each of the tested antibodies, regardless of species or antibody labels. In contrast, no binding was observed when testing GFP-Ft under the same conditions (data not shown). We can therefore safely assume that the hybrid protA-Ft can be equipped with any IgG as long as the antibody has sufficient affinity towards SpA.^49^

We demonstrated the performance of the activated hybrid protA-Ft by targeting the extracellular domain of the death receptor CD95. Here, protA-Ft’s binding capabilities only come in to play in the presence of an αCD95 antibody (see Figure 4A). Thus, prior to the microshower with protA-Ft, cells were incubated with respective antibody for 15 min. Afterwards, a microshower as described previously (Figure 3A) was applied for 15 min. While no binding was observed in cells not treated with antibody, a slight increase occurred when showering antibody-treated wild type cells with protA-Ft. This minimal increase might be caused by the intrinsic presence of natural CD95 receptor on the cell’s membrane.

**Figure 4.**
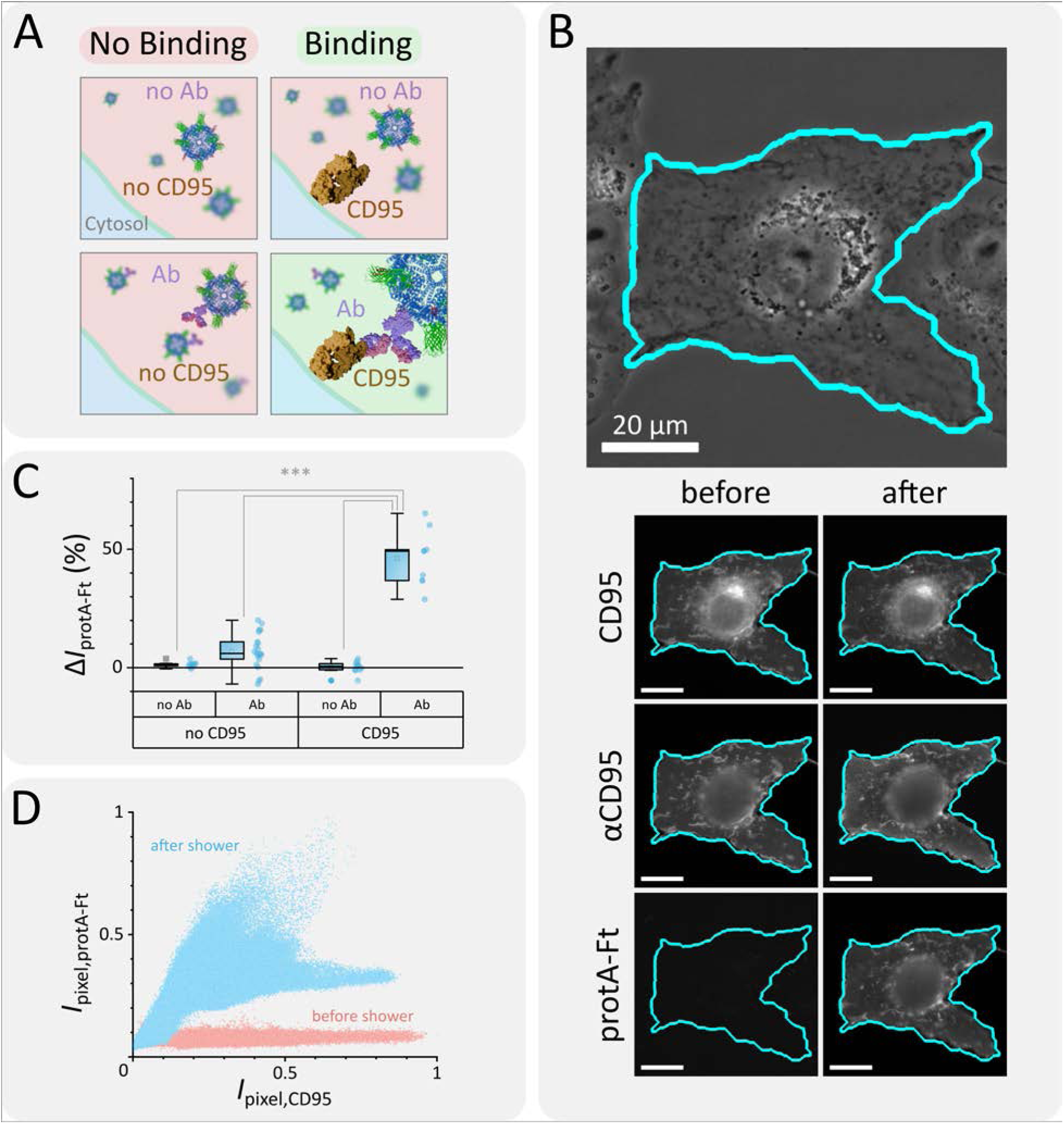
Targeting of CD95 using protA-Ft in combination with αCD95 antibody. (A) Schematic experimental setup. Cos7 cells were incubated with αCD95 for 15 min and subsequently showered with a microcapillary filled with protA-Ft for 15 min. Binding to cells occurs on cells that are overexpressing CD95 and only in the presence of the antibody. (B) Microscopy data of exemplary antibody-treated cell before and after protA-Ft microshower. Shown are phase contrast image, red channel (CD95 – mCherry, far-red channel (αCD95 – APC), and green channel (Ferritin – mEGFP). (C) Intensity increase in protA-Ft’s fluorescence channel after microshower for 15 min in wild type cells (no CD95) and TfR1-overexpressing cells (CD95) with and without previous antibody-treatment (Ab, no Ab). A one-way ANOVA was performed to test for differences between groups (*** = p < 0.001). (D) Scatter plot of pixel values from CD95 channel vs. protA-Ft channel.

However, treatment of CD95-overexpressing cells with antibody followed by protA-Ft led to a significantly larger rise of signal by 46 ± 12 % (see Figure 4B and C). Hence, it becomes apparent, that protA-Ft binding to CD95 is indeed enabled by the antibody. This is also confirmed by comparing pixel values of protA-Ft and CD95 (Figure 4D). As seen for the binding of protA-Ft to TfR1, the two intensities are linearly dependent and thus strongly co-localized. As for TfR1 experiments, no binding was observed with PEGylated protA-Ft, regardless of presence or absence of antibody (data not shown).

The key points of this work are the specific targeting of MRs and, if applicable, the subsequent activation, namely of CD95, by incubation with protA-Ft. It was shown in Figure 3 and Figure 4 that protA-Ft is able to specifically target both its natural receptors, demonstrated by the exemplary receptor TfR1, as well as other MRs through antibody-mediated binding. Figure 5 shows two examples for cells over-expressing TfR1 or CD95 respectively, before and at certain timepoints after microshower with protA-Ft. In TfR1 cells, the presence of protA-Ft leads to rapid formation of clusters of TfR1, illustrated by the transition of homogenous fluorescence signal to a granular image directly after the shower. Within 2 h, these clusters grow and steadily move towards the cell’s nucleus.

**Figure 5.**
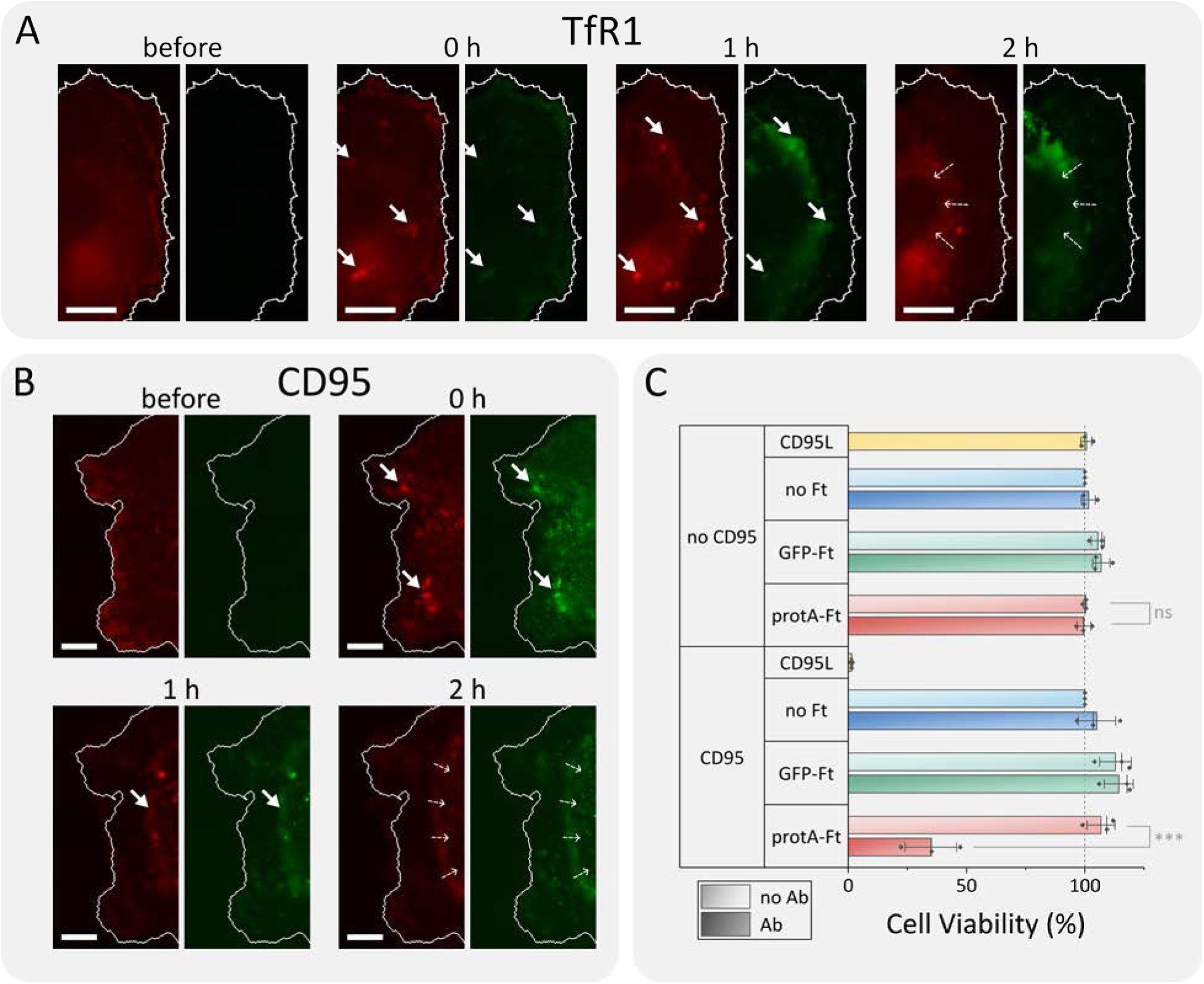
Exemplary Cos7 cells over-expressing (A) TfR1 (red), and (B) overexpressing CD95 (red) and treated with αCD95 antibody. Images were taken before or at denoted timepoints after a 15 min microshower with protA-Ft (green). Solid arrows are pointing to formed clusters. Dotted arrows indicate movement of protA-Ft and the respective receptor towards the cell nucleus. All scale bars are 10 µm. (C) CTB Viability assay of HeLa cells overexpressing CD95 (‘CD95’) or with CD95 being knocked out (‘no CD95’). Cells were incubated overnight with or without αCD95 antibody and with or without protA-Ft or GFP-Ft. CD95L served as negative control, that triggers apoptosis upon contact with CD95. A one-way ANOVA was performed to test for differences between groups (ns = not significant, *** = p < 0.001).

A similar behavior was observed on CD95 cells: immediately after protA-Ft initiation, clusters appeared in both the CD95 and protA-Ft channels. Rapidly after the microshower, a ring of CD95 and protA-Ft formed in the periphery of the cell that moved towards the nucleus within 1 h. Due to the highly synchronized and directed movement, it seems that endocytotic vesicles containing both protA-Ft as well as CD95 are being transported towards the center of the cell.

Probing of CD95 activation via protA-Ft incubation was performed using a cell viability assay. To this end, two cell lines were used – a knockout cell line without CD95 and a cell line stably overexpressing CD95. Both cell lines were incubated with different combinations of CD95-specific Ab, protA-Ft, GFP-Ft or no Ft. As a positive control, CD95L which induces apoptosis in the presence of CD95, was added to the cells. As shown in Figure 5B only the combination of CD95, protA-Ft and the CD95-specific Ab was able to induce apoptosis. Cell survival rates under these conditions dropped to 35 ± 13 %, while all other conditions remained at survival rates within the margin of error of the negative control. In the positive control with CD95L in the presence of CD95, 1.4 ± 0.5 % of all cells survived. This demonstrates, that only through the clustering of CD95 using our engineered protA-Ft, apoptosis can be initiated.

In this comprehensive study we assessed the potential of ferritin as a targetable NP. Due to its protein nature, stability and biocompatibility, ferritin serves as an excellent targeting system. By creating hybrid ferritin cages, multifunctionality can be readily introduced to the NP without the need for downstream chemical modifications. We demonstrated this by fusing mEGFP- and protA-labeled subunits that combined the capabilities of both modifications into one cage. The implementation of protA, a minimal binding model of the antibody-binding SpA, gave rise to a targeting system that can be easily adapted to experimental needs by simple means of incubation with antibodies. We demonstrated its targeting capabilities by targeting our hybrid protA-Ft to both TfR1 overexpressing cells via natural binding of ferritin and to CD95 via antibody-mediated binding, two receptors that are connected to cancer. Through both targeting routes, protA-Ft induced cluster formation of the targeted receptor. The intensive accumulation of CD95 through protA-Ft was enough to exceed cellular thresholds and ultimately induce apoptosis.

Conclusively, our manufactured hybrid protein cages are an easily operatable, highly flexible nanosystem for site-specific cellular targeting. Combination of this system with methods for loading small molecules into ferritin, such as anti-cancer drugs,^59–61^ could create a potent carrier for targeted drug delivery.^62^ Our cellular studies gave indications, that protA-Ft was internalized by the targeted cells, a useful property for drug delivery.^63^ Furthermore, by implementation of a magnetic core into the hybrid ferritin cage, a versatile tool for spatial manipulation and accumulation of MRs in living cells can be created.^17,52–54,64^ Such a tool would have the potential to explore previously unknown mechanisms underlying the signaling pathways of membrane receptors.

## ASSOCIATED CONTENT

Supporting Information: Detailed description of materials and methods and supplementary figures.

## AUTHOR INFORMATION

Author Contributions: The manuscript was written through contributions of all authors. All authors have given approval to the final version of the manuscript.

Conceptualization: CM Methodology: AN Software: AN Validation: AN, CM Formal analysis: AN

Investigation: AN, CS, LZ, ACF Resources: CM

Data Curation: AN Writing-original draft: AN

Writing-review & Editing: AN, CS, LZ, ACF, CM Visualization: AN

Supervision: CM

Project administration: CM Funding acquisition: CM

Notes: The authors declare no competing financial interest.

## ACKNOWLEDGMENT

CM acknowledges financial support by the DFG Collaborative Research Center 1208 “Identity and dynamics of biological membranes” (project ID 267205415), the DFG Collaborative Research Center 1535 ‘MiBiNet’ (project ID 458090666). CM and AN acknowledges financial support by the ‘Freigeist-fellowship’ of VolkswagenFoundation. The authors acknowledge the DFG and the State of North Rhine–Westphalia for funding the cryo-TEM (INST 208/749-1 FUGG) hosted by the Centre of Advanced Imaging (CAi, Heinrich-Heine University). We thank the Institute of Molecular Physical Chemistry (University of Düsseldorf) of Prof. Seidel for providing lab space and laboratory equipment, the Institute of Computational Pharmaceutical Chemistry (University of Düsseldorf) of Prof. Gohlke for providing the plate reader, the Institute of Synthetic Membrane Systems of Prof. Kedrov (University of Düsseldorf) for providing lab space and equipment, the Institute of Biochemistry I (University of Düsseldorf) of Prof. Schmitt for providing the cell disruptor, and the Institute of Macromolecular Chemistry of Prof. Hartmann (University of Düsseldorf) for providing the Zetasizer.

## ABBREVIATIONS

MR: membrane receptor

SpA: Protein A from S. aureus;

protA: engineered Z-domain of *S. aureus* Protein A

mEGFP: monomeric Enhanced Green Fluorescent Protein

Ft: Ferritin

protA-Ft: ferritin with mEGFP and protA subunits fused to its cage

GFP-Ft: ferritin with mEGFP subunits fused to its cage

HCF: Heavy Chain Ferritin

LCF: Light Chain Ferritin

GFP-HCF: HCF subunits fused to mEGFP

protA-HCF: protA subunit fused to HCF

TEM: Transmission Electron Microscopy

DLS: Dynamic Light Scattering

SDS-PAGE: Sodiumdodecly Sulphate-Polyacrylamide Gel Electrophoresis

