## Supplementary Information for "Semisynthetic ferritin nanocages for flexible, site-specific targeting, cluster-formation and activation of membrane receptors"

### Inhalt

### MATERIALS AND METHODS

#### Bacterial transformation and genetic engineering

As described in the main text, one ferritin cage consists of 24 subunits. For this study, cages only comprised of modified heavy chain ferritin (HCF; PDB code: 2FHA) subunits. Through genetic engineering two constructs were prepared: HCF with an N-terminal mEGFP (as a generous gift from the Coppey/Haji laboratory at the Laboratoire Physico-Chimie, Institut Curie, Paris, France and the Piehler laboratory at the University of Osnabrück, Germany), denoted GFP-HCF (see Figure S3) and HCF with an N-terminal protA (see Figure S4), a minimal binding domain of Protein A from *S. aureus*,[1–3] denoted protA-HCF. Additionally, GFP-HCF was equipped with a His-tag at its N terminus. cDNA of GFP-HCF was inserted into the vector pET21a (reporter gene: *ampR*), protA-HCF was inserted into pET24a (reporter gene: *kanR*).

#### Homopolymeric Ferritin

In order to produce Ferritin cages consisting of only one subunit, *E. coli* from the strain BL21-CodonPlus (DE3)-RIPL (Agilent, Santa Clara, CA, USA), were transformed with one of the two HCF constructs as described previously.[4]

#### Heteropolymeric Ferritin

A two-step bacterial transformation was performed to create a hybrid ferritin cage consisting of both GFP-HCF and protA-HCF. In the first step, bacteria were transformed as previously established.[4] After transformation, bacteria were plated on LB-agar plates containing the respective antibiotic and incubated at 37 °C overnight. The next day, a single colony was picked and transferred into 5 ml of LB containing 1 g/l of glucose for an overnight culture. Afterwards, 500 µl of the overnight culture were used to inoculate 50 ml of LB medium without antibiotics. The culture was grown up to an $OD_{600}$ of 0.4 to 0.5. Bacteria were then transferred into a centrifuge tube, chilled on ice for 15 min, and then centrifuged at 4 °C and 4,000 g for 10 min. The cell pellet was resuspended in ice-cold resuspension buffer (100 mM $CaCl_2$). After another 15 min on ice, bacteria were again centrifuged at 4 °C and 4,000 g for 10 min and collected in 2 ml of competence buffer (100 mM $CaCl_2$, 15 % v/v glycerol). As referenced earlier, the subsequent transformation was performed with 50 µl of these competent bacteria.[4] With this method, bacteria were first transformed with GFP-HCF and then, after rendering the transformed bacteria competent, with protA-HCF, to express both subunits simultaneously and produce hybrid ferritin cages.

#### Protein expression and purification

Production of protein and the subsequent purification were performed as referenced earlier.[4] Owing to the incorporated His-tag in both ferritin constructs, after heat denaturation of unwanted proteins and the subsequent centrifugation, the protein solution was purified using immobilized metal affinity chromatography (IMAC) in an NGC system (Bio-Rad, Hercules, CA, USA). The system – together with a $Ni^{2+}$ activated HisTrap HP column (Cytiva, Marlborough, MA, USA) – was flushed with binding buffer BB (20 mM HEPES, 100 mM

NaCl, 20 mM imidazole, pH 8.0, filtered through 0.2 µm mesh, degassed) and elution buffer EB (20 mM HEPES, 100 mM NaCl, 300 mM imidazole, pH 8.0, filtered through 0.2 µm mesh, degassed) followed by another flush with BB. In preparation for IMAC, 20 mM imidazole was added to the sample solution. The sample was then loaded onto the column and washed with BB, to remove unspecifically bound protein. The target protein was eluted by applying an elution gradient from 0 to 100 % of EB over 10 column volumes (CV). Eluate was collected with a fraction size of 3 ml. Fractions containing the target protein were combined, reconcentrated, washed with HEPES buffer (20 mM HEPES, 100 mM NaCl, pH 8.0, filtered through 0.2 µm mesh, degassed), and aliquoted for further needs.

Transmission electron microscopy (TEM)

For TEM analysis, samples were diluted to 0.05 mg/ml and a droplet of 7 µl was placed on a Ni grid with Formvar coating (S162N, PLANO GmbH, Wetzlar, Germany). Proteins were left to sediment for 1 min and then removed from the grid using filter paper. The grid was then dipped for 3 s into a droplet of uranyl acetate solution (2 %), dried with filter paper and placed on top of a second drop of uranyl acetate. After 30 s, the grid was removed from the drop, dried using filter paper, and left to dry for 15 min.

Samples were imaged using a Jeol JEM-2100Plus (Akishima, Tokyo, Japan) at an acceleration voltage of 80 kV. Images were analyzed using Gatan Micrograph Suite (Gatan Inc., Pleasanton, CA, USA) and Matlab 2023a (Mathworks, Natick, MA, USA).

Fluorescence linked immunosorbent assay (FLISA)

The binding behavior between the ferritin constructs and target antibodies was assessed using FLISA. For this, 50 µl of antibody solution (1 µg/ml) were added into the wells of a 96-well plate (half area, black, medium binding, REF 675076, Greiner Bio-One, Kremsmünster, Austria) and incubated for 1 h at room temperature. The analyzed antibodies were a human IgG1 control antibody (REA Control Antibody, human IgG1, pure, 130-129-977, Miltenyi Biotec, Bergisch Gladbach, Germany), a murine IgG2a (Art. No. 130-106-546, Miltenyi Biotec), and an APC-labeled human IgG1 (Art. No. 130-113-446, Miltenyi Biotec). Wells were then filled to their maximum by adding 130 µl of blocking buffer (20 mM HEPES, 100 mM NaCl, pH 8.0, 2 % (w/v) BSA) and incubated overnight at 4 °C. The next day, serial dilutions of ferritin samples were prepared freshly. Afterwards, wells were emptied and washed three times with 180 µl of HEPES-T (20 mM HEPES, 100 mM NaCl, pH 8.0, 0.05 % (v/v) Tween20). After the last washing step, it is highly important to remove all traces of liquid from each well. Then, 50 µl of the ferritin constructs' dilution series were added to each corresponding well and incubated at room temperature for 1 h in the dark. Wells were then washed again three times and afterwards filled with 50 µl of HEPES buffer (20 mM HEPES, 100 mM NaCl, pH 8.0).

The FLISA was analyzed using a Tecan Infinite M Plex (Tecan Group, Männedorf, Switzerland) in fluorescence mode (Excitation: 488 nm | Emission: 540 nm). The obtained data was analyzed using Matlab. Normalized data was fitted using four-parameter logistic regression:

$$I(x) = D + \frac{A - D}{1 + \left(\frac{x}{C}\right)^B} \ ,$$

with the plateau at infinite concentration *A*, the asymptote at zero concentration *D*, the slope parameter *B* and the point of inflection – or $K_D$ value – *C*. The fit was forced to stay within the Interval of 0 to 1, since the data sets were background subtracted and normalized to the positive control (ferritin incubated directly in the well).

Dynamic light scattering (DLS)

DLS measurements were performed using a Zetasizer Nano ZS (Malvern Panalytical Ltd, Malvern, UK). Prior to measurement, all samples were centrifuged at 10,000 g for 10 min and only the supernatant was analyzed. 20 µl of sample solution at a concentration between 0.1 and 1.0 mg/ml was placed into a Ultra-Micro-cuvette, placed in the Zetasizer, and left to equilibrate to 25°C for 3 min. The sample was analyzed in five measurement cycles, each containing 15 runs of 10 s. The delay between each measurement cycle was 1 min. The measurement angle was set to 173° backscatter. The obtained data was analyzed according to the number distribution in order to realistically weight the measured nanoparticles.

Spectroscopy and degree of labeling

In order to determine the concentration and degree of labeling (DoL; number of fluorophores per nanoparticle) of our ferritin constructs, spectroscopical analyses were conducted. In this work, it was determined spectroscopically using a NanoPhotometer NP80 (Implen, Munich, Germany). The absorbance of samples was measured in the range of 200 to 900 nm and the mean of 550 to 900 nm was subtracted from each curve as background. As described in the main text, Lambert-Beer’s law was used for the analysis:

$$DoL = \frac{\varepsilon_{488,GFP}}{\varepsilon_{280,Ft}} \frac{A_{488}}{(A_{280} - A_{488} \cdot CF_{GFP})},$$

with the following parameters: $\varepsilon_{488,GFP}$ = 56,000 M-1cm-1, $\varepsilon_{280,Ft}$ = 462,860 M-1cm-1, $A_{280}$ and $A_{488}$ as measured absorbances of the sample at wavelength 280 and 488 nm respectively, and the correction factor $CF_{GFP}$ as the ratio of $A_{280}$ over $A_{488}$. This correction factor was determined from a separately, pure sample of mEGFP to be 0.54.

SDS-PAGE

Protein samples were mixed in a 1:1 ratio with 2x Laemmli buffer (Cat. No. 42526.01, Serva, Heidelberg, Germany) denatured for 5 min at 100 °C and centrifuged for 15 min at 4 °C and 15,000 g. The supernatant was loaded onto a self-cast 11% SDS gel and run at 100 V for 90 to 120 min. Afterwards, the gel was washed with water several times and then stained using Coomassie blue stain (SimplyBlue™ SafeStain, Thermo Fisher Scientific, Waltham, MA, USA) for 1 h on a horizontal shaker. After staining, the gel was washed with water and destained for 1 h in water, also shaking. Gels were then imaged using a GelDoc Go Imaging System (Bio-Rad).

The images were processed in Fiji.[5] After background subtraction (rolling ball radius: 50 px, sliding paraboloid) Fiji's gel analysis tool was used to plot band intensities. The area under the peaks was used for estimations of protein contents.

### Cell culture and transient transfection

Adherent Cos7 cells, HeLa CD95-Knockout cells, and HeLa overexpressing CD95 were cultivated at 37 °C in a 5 % $CO_2$ atmosphere. Cells were cultured in Dulbecco's Modified Eagle Medium (DMEM) supplemented with 10 v% fetal calve serum (FCS) and 1 v% PenStrep (all from Thermo Fisher Scientific). Cos7 cells were seeded onto imaging dishes 24 h before transfection. Transfection of Cos7 was carried out the day before imaging using the transfection agent ViaFect (Promega, Madison, Wisconsin, USA) according to the manufacturer's protocol. During imaging, cells were kept in Leibovitz's L-15 Medium (L-15) without phenol red. HeLa cells for CTB assay were seeded in a 96-well plate 24 h before the experiment.

### Microscopy and microshower

Images were taken with an inverted epifluorescence microscope (IX83 by Olympus, Shinjuku, Tokyo, Japan) in combination with a 60x oil objective (N.A. = 1.25, PH3, Olympus UPLFLN60XOIPH/1,25, Olympus). Microshower of protA-Ft was performed using a microinjection setup, consisting of a micromanipulator (InjectMan 4, Eppendorf, Hamburg, Germany), a pump system (FemtoJet 4i, Eppendorf), and matching microcapillaries (Femtotip II, inner diameter of 500 nm, Eppendorf). The loaded ferritin solution was at a concentration of 1 mg/ml and centrifuged for 10 min at 10,000 g to remove any agglomerates before loading into the capillary. The pump system's compensation pressure was adjusted to sustain a constant flow out of the microcapillary during the microshower. For CD95 experiments, cells were incubated for 15 min with CD95-specific antibody (Art. No. 130-108-066, Miltenyi Biotec) at a dilution of 1:50 in L-15 medium. Then, cells were washed with Dulbecco's Phosphate Buffered Saline PBS (DPBS, Thermo Fisher Scientific) and imaged in L-15 medium.

### CellTiter-blue® viability assay

Apoptosis evaluation was performed using CellTiter®-blue (CTB) assay (Promega, Fitchburg, WI, USA) according to the manufacturer's manual and as described previously.[6] In short, HeLa CD95-Knockout and HeLa stably overexpressing CD95 were seeded into 96-well plates, with 25,000 cells per well, the day before. Cells were washed with DPBS, once and incubated in DMEM and the respective mixture of CD95-specific antibody (Miltenyi), protA-Ft, GFP-Ft, and CD95 ligand (CD95L), all at concentration of 1 µg/ml in DMEM. After 16 h, cells were washed once and incubated for 3 h in a 1:10 mix of DMEM and CTB solution. Well plates were analyzed using Infinite M Plex plate reader (Tecan, Männedorf, Switzerland) with 560 nm excitation and 590 nm emission over 25 flashes and 5 repetitions. Three biological replicates were analyzed. Data was background subtracted (control well with all cells killed using 0.1% of Triton X-100 (PanReac AppliChem, Darmstadt, Germany) for 30 min, as 0 % cell viability) and normalized to the positive control (cells only incubated with DMEM, as 100 % cell viability).

SUPPLEMENTARY FIGURES

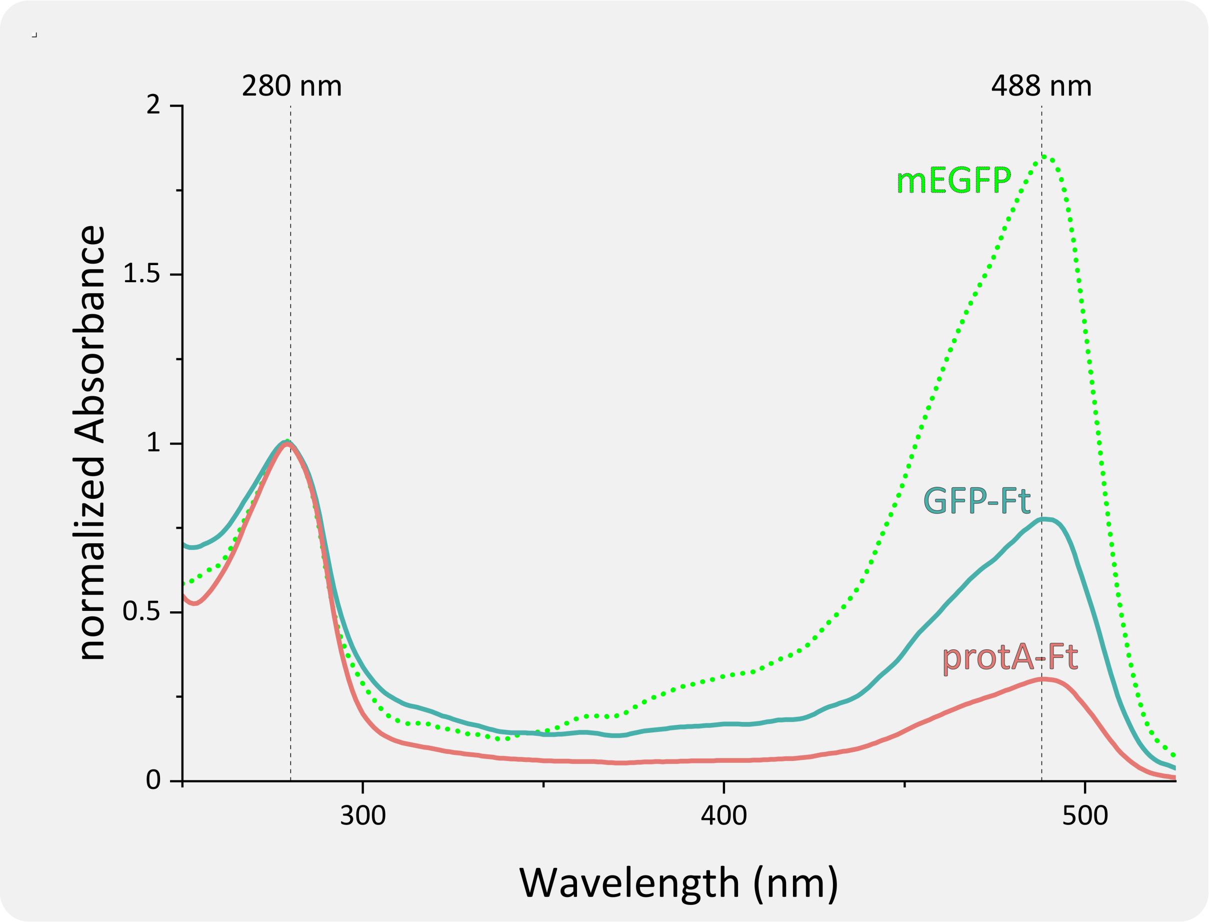

*Figure S1 – Absorption spectra of pure GFP-Ft and protA-Ft for determination of the degree of labeling (DoL). Pure mEGFP was measured as a reference for the calculation of the correction factor of mEGFP. All spectra are normalized to the absorbance at 280 nm.*

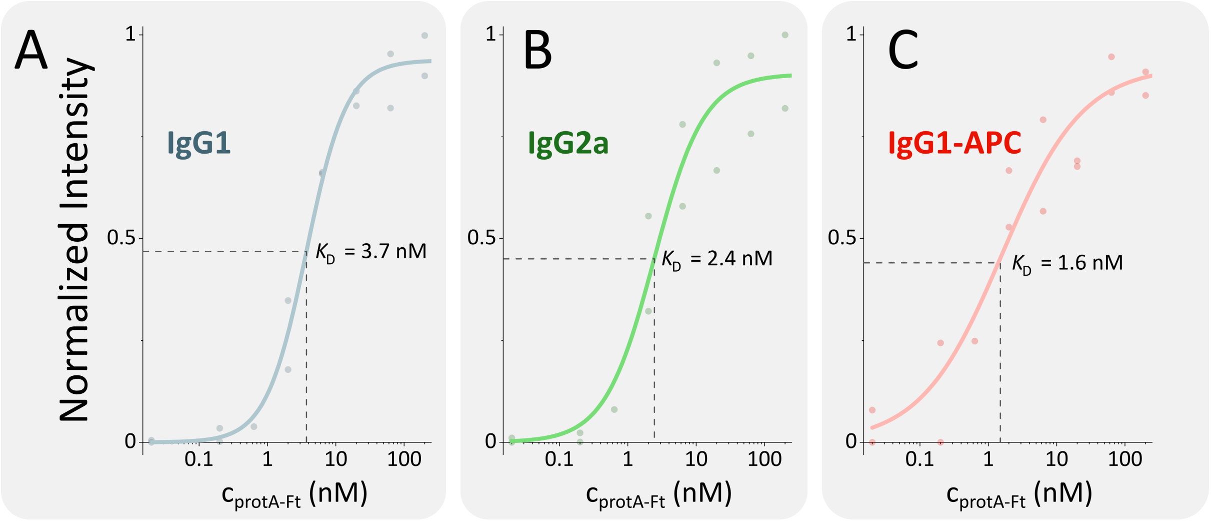

*Figure S2 – Results of FLISA with three different antibodies and incubation with protA-Ft in different concentrations. The $K_D$ was obtained by fitting a 4PL curve (see Materials and Methods) to the data. Data were obtained as three technical replicates.*

Figure S3 – Sequence of His6–mEGFP–Linker–HCF

His6

mEGFP

Linker

HCF

DNA Sequence

ATGCATCATCATCACCATCACGGCAGCGGC-
GTGAGCAAGGGCGAGGAGCTGTTCACCGGGGTGGTGCCCATCCTGGTCGAGCTGGACGGCGACGTAAACGG
CCACAAGTTCAGCGTGTCCGGCGAGGGCGAGGGCGATGCCACCTACGGCAAGCTGACCCTGAAGTTCATCTG
CACCACCGGCAAGCTGCCCGTGCCCTGGCCCACCCTCGTGACCACCCTGACCTACGGCGTGCAGTGCTTCAGCC
GCTACCCCGACCACATGAAGCAGCACGACTTCTTCAAGTCCGCCATGCCCGAAGGCTACGTCCAGGAGCGCAC
CATCTTCTTCAAGGACGACGGCAACTACAAGACCCGCGCCGAGGTGAAGTTCGAGGGCGACACCCTGGTGAA
CCGCATCGAGCTGAAGGGCATCGACTTCAAGGAGGACGGCAACATCCTGGGGCACAAGCTGGAGTACAACTA
CAACAGCCACAACGTCTATATCATGGCCGACAAGCAGAAGAACGGCATCAAGGTGAACTTCAAGATCCGCCAC
AACATCGAGGACGGCAGCGTGCAGCTCGCCGACCACTACCAGCAGAACACCCCCATCGGCGACGGCCCCGTG
CTGCTGCCCGACAACCACTACCTGAGCACCCAGTCCAAGCTGAGCAAAGACCCCAACGAGAAGCGCGATCACA
TGGTCCTGCTGGAGTTCGTGACCGCCGCCGGGATCACTCTCGGCATGGACGAGCTGTACAAG-
GGATCCGGTGCAGAATTC-
ATGACGACCGCGTCCACCTCGCAGGTGCGCCAGAACTACCACCAGGACTCAGAGGCCGCCATCAACCGCCAGA
TCAACCTGGAGCTCTACGCCTCCTACGTTTACCTGTCCATGTCTTACTACTTTGACCGCGATGATGTGGCTTTGA
AGAACTTTGCCAAATACTTTCTTCACCAATCTCATGAGGAGAGGGAACATGCTGAGAAACTGATGAAGCTGCA
GAACCAACGAGGTGGCCGAATCTTCCTTCAGGATATCAAGAAACCAGACTGTGATGACTGGGAGAGCGGGCT
GAATGCAATGGAGTGTGCATTACATTTGGAAAAAAATGTGAATCAGTCACTACTGGAACTGCACAAACTGGCC
ACTGACAAAAATGACCCCCATTTGTGTGACTTCATTGAGACACATTACCTGAATGAGCAGGTGAAAGCCATCAA
AGAATTGGGTGACCACGTGACCAACTTGCGCAAGATGGGAGCGCCCGAATCTGGCTTGGCGGAATATCTCTTT
GACAAGCACACCCTGGGAGACAGTGATAATGAAAGCTAG

Protein Sequence

MHHHHHHGSG-
VSKGEELFTGVVPILVELDGDVNGHKFSVSGEGEGDATYGKLTLKFICTTGKLPVPWPTLVTTLTYGVQCFSRYPDH
MKQHDFFKSAMPEGYVQERTIFFKDDGNYKTRAEVKFEGDTLVNRIELKGIDFKEDGNILGHKLEYNYNSHNVYIM
ADKQKNGIKVNFKIRHNIEDGSVQLADHYQQNTPIGDGPVLLPDNHYLSTQSKLSKDPNEKRDHMVLLEFVTAAGI
TLGMDELYK-GSGAEF-
MTTASTSQVRQNYHQDSEAAINRQINLELYASYVYLSMSYYFDRDDVALKNFAKYFLHQSHEEREHAEKLMKLQN
QRGGRIFLQDIKKPDCDDWESGLNAMECALHLEKNVNQSLLELHKLATDKNDPHLCDFIETHYLNEQVKAIKELGD
HVTNLRKMGAPESGLAEYLFDKHTLGDSDNES

Figure S4 – Sequence of protA–Linker–HCF

protA

Linker

HCF

DNA Sequence

ATGGTTGACAACAAATTCAACAAAGAACAGCAGAACGCGTTCTACGAAATCCTGCACCTGCCGAACCTGAACG
AAGAACAGCGTAACGCGTTCATCCAGTCTCTGAAAGACGACCCGTCTCAGTCTGCGAACCTGCTGGCGGAAGC
GAAAAAACTGAACGACGCGCAGGCGCCGAAA-GGATCCGGTGCAGAATTC-
ATGACGACCGCGTCCACCTCGCAGGTGCGCCAGAACTACCACCAGGACTCAGAGGCCGCCATCAACCGCCAGA
TCAACCTGGAGCTCTACGCCTCCTACGTTTACCTGTCCATGTCTTACTACTTTGACCGCGATGATGTGGCTTTGA
AGAACTTTGCCAAATACTTTCTTCACCAATCTCATGAGGAGAGGGAACATGCTGAGAAACTGATGAAGCTGCA
GAACCAACGAGGTGGCCGAATCTTCCTTCAGGATATCAAGAAACCAGACTGTGATGACTGGGAGAGCGGGCT
GAATGCAATGGAGTGTGCATTACATTTGGAAAAAAATGTGAATCAGTCACTACTGGAACTGCACAAACTGGCC
ACTGACAAAAATGACCCCCATTTGTGTGACTTCATTGAGACACATTACCTGAATGAGCAGGTGAAAGCCATCAA
AGAATTGGGTGACCACGTGACCAACTTGCGCAAGATGGGAGCGCCCGAATCTGGCTTGGCGGAATATCTCTTT
GACAAGCACACCCTGGGAGACAGTGATAATGAAAGCTAG

Protein Sequence

MVDNKFNKEQQNAFYEILHLPNLNEEQRNAFIQSLKDDPSQSANLLAEAKKLNDAQAPK-GSGAEF-
MTTASTSQVRQNYHQDSEAAINRQINLELYASYVYLSMSYYFDRDDVALKNFAKYFLHQSHEEREHAEKLMKLQN
QRGGRIFLQDIKKPDCDDWESGLNAMECALHLEKNVNQSLLELHKLATDKNDPHLCDFIETHYLNEQVKAIKELGD
HVTNLRKMGAPESGLAEYLFDKHTLGDSDNES*
